# Canopy health, but not *Candidatus* Liberibacter asiaticus Ct values, are correlated with fruit yield in Huanglongbing affected sweet orange trees

**DOI:** 10.1101/2021.09.07.459315

**Authors:** Amit Levy, Taylor Livingston, Chunxia Wang, Diann Achor, Tripti Vashisth

**Author notes:** **Corresponding authors:** Tripti Vashisth; Amit Levy. **Author Contributions:** AL and TV designed the experiments.TL, CW and DA conducted the work. AL, CW and TV wrote the manuscript.

## Abstract

In Florida, almost all citrus trees are infected with Huanglongbing (HLB), caused by the gram-negative, intracellular phloem limited bacteria *Candidatus* liberibacter asiaticus (CLas). Distinguishing between the severely and mildly sick trees is important for managing the groves and testing new HLB therapies. A mildly sick tree is one that produces higher fruit yield, compared to a severely sick tree, but measuring yields is laborious and time consuming. Here we characterized HLB affected sweet orange trees in the field in order to identify the specific traits that are correlated with the yields. We found that canopy volume, fruit detachment force (FDF) and the percentage of photosynthetically active radiation interception in the canopy (%INT) were positively correlated with fruit yields. Specifically, %INT measurements accurately distinguished between mild and severe trees in independent field trials. We could not find a difference in the Ct value between high and low producing HLB trees. Moreover, Ct values did not always agree with the number of CLas in the phloem that were visualized by transmission electron microscopy. Overall, our work identified an efficient way to distinguish between severe and mild HLB trees in Florida by measuring %INT and suggests that health of the canopy is more important for yields than the Ct value.

## Introduction

Huanglongbing (HLB), or citrus greening, has devastated citrus production in Florida and is spreading worldwide. The disease is caused by the gram-negative, intracellular phloem limited bacteria *Candidatus* liberibacter asiaticus (CLas) and is transmitted by the Asian citrus psyllid (ACP; *Diaphorina citri*). HLB results in bitter and asymmetrical fruit, seed abortion and premature fruit drop [1]. Other disease symptoms include blotchy mottled (non-symmetrical chlorosis or mottling), pale yellow leaves, yellow shoots, corky veins, stunting, damaged roots, and twig dieback. In leaves and stems, a microscopic phenotype typically associated with CLas infection is the accumulation of callose inside the phloem sieve plate pores [2, 3], which begins at early stages of the disease and leads to the collapse of the phloem at more advanced disease stages. As a result, the products of photosynthesis cannot be effectively transported from the leaves, so they are converted into starch and deposited among the grana stacks of the chloroplast [3]. Eventually the starch deposits become so large that they destroy the grana stacks, rendering the chloroplast inoperative [2].

In Florida, HLB reduced the production of citrus by 74% [4]. Current management options for HLB are limited and heavily rely on the application of insecticides for controlling the insect vector, *D. citri*. However, insecticide treatment is unsustainable due to the development of resistance among ACP populations, damage to the environment and beneficial organisms, and high cost [1]. Similar challenges and problems also arise when applying antibiotics to control the bacterial population in the trees. Enhanced nutritional treatments, such as controlled release fertilizer (CRF), were shown to increase fruit yield and quality [5], but these treatments can take long time to show impact on yield. There is an ongoing effort to develop HLB resistant or tolerant citrus varieties, but currently there is no known such variety that can be used in the field.

The successful development of HLB therapies requires an efficient method to test the effectiveness of the treatments in improving yield and tree health. HLB has spread to all growing areas in Florida, and there are no uninfected control trees. There is a need for a method that can easily determine the impact of treatments on HLB severity. The most common method to determine HLB levels is by quantifying CLas with polymerase chain reaction (PCR). This method is used to determine if treatments result in lower amount of CLas in the plants [6–9]. Regular PCR is used in order to detect CLas DNA in the plant sample, and quantitative PCR (qPCR) is used to measure the number of copies of CLas DNA, corresponding to its titer [10]. With qPCR, the impacts of different treatments on HLB is usually determined by the Ct values [11, 12], and the relative change in phytopathogen titer in response to treatment is determined with the 2^−ΔΔCt^ method [13]. By including known amounts of plasmid with the corresponding CLas sequence in the qPCR analysis, the number of cells of CLas per gram of plant tissue sample can be also predicted [14]. These values are then used to quantify the relative change in CLas population in response to treatment [6]. Other techniques include microscopic, spectroscopic and imaging techniques, but those are more complex and expensive [15]. In the field, another technique to determine HLB levels is Disease Index Rating [7], however it’s a non-standardized, subjective rating method.

For citrus growers, the most important and relevant value for scoring HLB mitigation treatments is its effect on the fruit yield of the tree. However, it can take multiple years to see an effect of treatment on the yield [16], obtaining this value is very laborious, and can only be done in certain time of the year. Moreover, in many cases, there is a need to determine the effect of treatments while trees are still in their juvenile state, for example when analyzing genetically edited or modified plants. Therefore, finding an easier and more available method to score HLB, that correlates with the fruit yield, is highly desired. Here, we tested various properties of HLB trees in the field, and determined which values correlated with the yield. Among others, we also tested if Ct value can be used to determine HLB status in the field. Our results show that fruit detachment force, canopy volume and photosynthetically active radiation interception in the canopy are all correlated to yield and can be used to determine the HLB disease status in the field. However, there was no correlation between yield and Ct value, therefore Ct value is not an appropriate measure for HLB. Moreover, our results suggest that Ct value does not necessarily provide an accurate estimate of the number of viable CLas cells.

## Materials and Methods

### Plant materials

Three distinct experimental orchards at the University of Florida Citrus Research and Education Center in Lake Alfred were used for this research. Orchard 1 and 2 included eighteen-year-old and eleven-year-old ‘Valencia’ sweet orange trees on Swingle citrumelo (Citrus paradisi × Poncirus trifoliata) rootstock, respectively; orchard 3 included nineteen-year-old ‘Hamlin’ sweet orange trees on Swingle citrumelo. Currently, because of the prevalence of HLB in Florida, it is unlikely to find healthy CLas-negative sweet orange trees as controls in open-air groves. Therefore, trees (at least 5 single tree replicates were used for each orchard) exhibiting mild and severe visual symptoms of HLB were selected. CLas titer, canopy volume, leaf area index (LAI), photosynthetically active radiation (PAR) interception percent in the canopy (%INT), root density, leaf nutrient, leaf SPAD index, disease index rating, fruit size, fruit detachment force (FDF), and yield were collected on individual trees on the day of fruit harvest. Comparing transmission electron microscopy (TEM) and Ct values was performed on 5-month-old seedlings of Citrus macrophylla and ‘Duncan’ grapefruit (C. paradisi) grown under greenhouse conditions at the Citrus Research and Education Center (Lake Alfred, FL).

### Genomic DNA extraction and *Ca.* L. asiaticus’ titers

Thirty random, fully mature leaves, free of any defect were collected. The midribs of these leaves were excised using a clean razor blade and stored at −80 °C until DNA extraction. Samples of the leaf tissue midrib (100 mg) were grinded by TissueLyser II (Qiagen, Germantown, MD 20874 USA) using liquid nitrogen, extracted in 0.5 ml of extraction buffer (100 mM Tris-HCL, pH8.0; 50 mM EDTA; 500 mM NaCl; 2% CTAB, and 2% PVP 40). The extract was incubated at 65°C for 30 min. Following incubation, 500 μl of 5 M potassium acetate was added, mixed thoroughly, and incubated on ice for 20 min. The mixture was centrifuged at 13,000 rpm for 10 min, 400 μl of supernatant was recovered, and DNA was precipitated by adding an equal volume of isopropanol and held at –20°C overnight. The DNA was pelleted, washed, and resuspended in 100 μl of water for PCR analysis. qPCR was performed with primers and probes (HLBas, HLBr, and HLBp) for the ‘*Ca.* L.asiaticus’ bacterium [17]) using ABI PRISM 7500 sequence detection system (Applied Biosystems, Foster City, CA) in a 10-μl reaction volume consisting of the following reagents: 200 nM (each) target primer (HLBas and HLBr), 100 nM target probe (HLBp), and 1× TaqMan qPCR Mix (Applied Biosystems). The amplification protocol was 94°C for 5 minutes followed by 40 cycles at 94°C for10 s and 58°C for 40 s. All reactions were performed in triplicate and each run contained one negative (DNA from healthy plant) and one positive (DNA from ‘*Ca.* L. asiaticus’-infected plant) control. Data were analyzed using the ABI 7500 Fast Real-Time PCR System with SDS software. The resulting cycle threshold (Ct) values were converted to the estimated bacterial titers using the grand universal equation *Y* = 13.82 –0.2866*X*, where *Y* is the estimated log concentration of templates and *X* is the qPCR Ct values, as described by Li et al. [18].

### Disease index rating

Each tree was rated based on the disease index rating system developed by the Citrus Research and Development Foundation in Lake Alfred, FL (https://crec.ifas.ufl.edu/extension/trade_journals/2016/2016_August_hlb.pdf). Briefly, each side of the tree was divided in quarters and was given a score out of 5 based on the HLB symptoms, where 0 means highly symptomatic and 5 means no HLB symptoms. All the scores were summed up to give the tree a total score out of 40.

### Canopy measurements

Canopy volume was at time of harvest for each tree individually. Canopy volume, expressed as cubic meters was calculated using a geometric prolate spheroid formula:

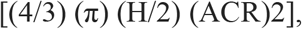

where π = 3.14, H = tree height, and ACR = average canopy radius. ACR was calculated by dividing tree diameter by 2 and calculating average radius. Tree diameter was measured in two directions: east to west (D1) and north to south (D2). Percentage increase in canopy volume was calculated as the increase in canopy volume from the time of pruning and at the end of the experiment.

HLB-affected trees often have sparse canopies; therefore, canopy density can be a good indicator of tree health and growth. Leaf area index (LAI) was measured for each tree to estimate canopy density. A plant canopy imager (CI-110; CID Bio-Science, Camas, WA) was used to measure the LAI and all the measurements were taken in the morning of a sunny day at the center of the canopy. The instrument was equipped with numerous light sensors to measure photosynthetically active radiation and with a global positioning unit to calculate the zenith angle for an accurate measurement of LAI. In addition to LAI, PAR interception (%INT) by the canopy was measured to provide another estimation of canopy density. Since PAR can be easily measured by quantum sensors, it is likely to be easily adopted by citrus growers. For %INT, the PAR was measured outside the canopy and inside the canopy (average of the PAR in each quarter of the canopy). The %INT was calculated by:

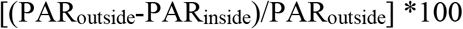

### Root density

Root density was measured at the time of harvest. From each tree, eight evenly spaced soil cores, ≈1 m from the trunk of the tree, were taken from the wetted zone. Roots were sifted, washed, and dried in an oven at 50 °C for 48 h, then the dry weight of the root samples was obtained. Root density was expressed as milligram of dry weight per cubic centimeter of soil.

### Leaf nutrient analysis

Thirty random leaves with intact petioles per tree were collected from nonfruiting branches. The collected leaves were washed using acidic soap, then the leaves were dried for 48 h in a convection oven (Thermo Fisher Scientific, Waltham, MA) and ground to a fine powder. Ground leaves were sent to Waters Agricultural Laboratories (Camilla, GA) to perform a standard leaf nutrient analysis.

### SPAD index

The soil plant analysis development (SPAD) index was used to estimate the total leaf chlorophyll content of thirty fully expanded, randomly selected, mature leaves per tree using a chlorophyll meter (MC-100 Chlorophyll Concentration Meter; Apogee Instruments, Logan, UT). The thirty SPAD values of each tree were averaged for a representation of the chlorophyll content of the tree.

### Yield, fruit size and fruit detachment force (FDF)

When the total soluble solids to titratable acidity ratio of fruit reached commercial harvest standards, the fruit were hand harvested by commercially trained harvesters. ‘Valencia’ trees in orchard 1 and 2 March 3 and 12, 2020, respectively and ‘Hamlin’ were harvested on December 4, 2019. Fruit yield is expressed as total kg of fruit per tree. For each tree, at time of harvest, 20 fruit attached to the tree branches were collected by clipping the branches. Only the branches bearing single fruit at the distal position were collected. All 20 fruits were used for fruit size measurement using a handheld digital caliper to measure the equatorial length on all 20 fruits. For each fruit, FDF was measured as well. FDF determines how much force is required for the fruit to be “pulled” or detached from the tree/branch, that was measured using a digital force gauge (Force One; Wagner Instruments, Greenwich, CT). FDF >6 kgf [19] is considered as a fruit that is not physiologically ready to abscise, FDF ≤6 kgf represents a fruit that has high tendency to drop at the time of collection.

### Transmission Electron Microscopy (TEM)

To compare CLas seen in TEM with Ct values, one half of a leaf midrib was subjected to CLas detection with qPCR, as described above, and the other half of the same midrib was subjected to TEM analysis to identify CLas in the phloem, performed as described by Achor et al. [20]. Briefly, Samples were fixed with 3% (v/v) glutaraldehyde in 0.1 M of potassium phosphate buffer at pH 7.2 for 4 h at room temperature, washed in phosphate buffer, then postfixed in 2% osmium tetroxide (w/v) in the same buffer for 4 h at room temperature. The samples were further washed in the phosphate buffer, dehydrated in a 10% acetone (v/v) series (10 min per step), and infiltrated and embedded in Spurr’s resin over 3 d. Sections (100-nm) were mounted on 200-mesh formvar-coated copper grids, stained with 2% aq uranyl acetate (w/v) and Reynolds lead citrate, and examined with a Morgagni 268 transmission electron microscope (FEI).

## Results

### Fruit yield correlates with several factors

Fruit yield is a major concern for citrus growers. In this study, we investigated different parameters, to find which ones are correlated with citrus yield in the field. The main parameters are listed in Table 1, which includes canopy volume, HLB disease index, root density, SPAD, LAI, %INT, fruit size, FDF, yield, Ct value and CLas population. Three different locations with two citrus varieties (Valencia on Swingle, and Hamlin on Swingle) were selected for the study, the results demonstrated that different factors influenced each other. As shown in Table 1, disease index had significantly negative correlation with root density, SPAD, LAI, %INT and FDF. FDF was positively correlated with root density, LAI, fruit size and %INT. Remarkably, we found that high yields in the field are correlated not only with high canopy volume but also with high %INT and FDF (with r = 0.542, 0.427, 0.468 respectively, *p*<0.05). The Ct value and CLas population were not correlated with yield, and only showed significantly positive correlation with canopy volume (Table 1).

**Table 1.** Correlation between different HLB diseases tree characteristics. Plain numbers are correlation coefficient (r) values and italicized numbers below are *p* values. Bold numbers indicate correlation with fruit yield. #-no statistically significant correlation was found.

|  | Canopy volume (m<sup>3</sup>) | Disease index | Root density (g/L) | SPAD | Leaf area index | %INT | Fruit size (mm) | Fruit detachment force (Kgf) | Yield (kg/tree) | Ct value of qPCR | Clas population /g of fresh tissues |
| --- | --- | --- | --- | --- | --- | --- | --- | --- | --- | --- | --- |
| Canopy volume (m<sup>3</sup>) | 1 | # | # | # | # | # | # | # | # | # | # |
| Disease index | # | 1 | # | # | # | # | # | # | # | # | # |
| Root density (g/L) | # | -0.456<br>*0.015* | 1 | # | # | # | # | # | # | # | # |
| SPAD | -0.402<br>*0.034* | -0.464<br>*0.013* | # | 1 | # | # | # | # | # | # | # |
| Leaf area index | -0.518<br>*0.005* | -0.404<br>*0.033* | 0.483<br>*0.009* | 0.476<br>*0.010* | 1 | # | # | # | # | # | # |
| %INT | # | -0.678<br>*0.000* | 0.412<br>*0.029* | # | 0.424<br>*0.024* | 1 | # | # | # | # | # |
| Fruit size (mm) | # | # | # | # | # | 0.695<br>*0.000* | 1 | # | # | # | # |
| Fruit detachment force (Kgf) | # | -0.547<br>*0.013* | 0.465<br>*0.039* | # | 0.464<br>*0.040* | 0.738<br>*0.000* | 0.615<br>*0.004* | 1 | # | # | # |
| Yield (kg/tree) | **0.542**<br>***0.003*** | # | # | # | # | **0.427**<br>***0.024*** | # | **0.468**<br>***0.038*** | 1 | # | # |
| Ct value of qPCR | 0.389<br>*0.041* | # | # | # | # | # | # | # | # | 1 | # |
| Clas population /g of fresh tissues | # | # | # | # | # | # | # | # | # | # | 0 |

**Table 2:**
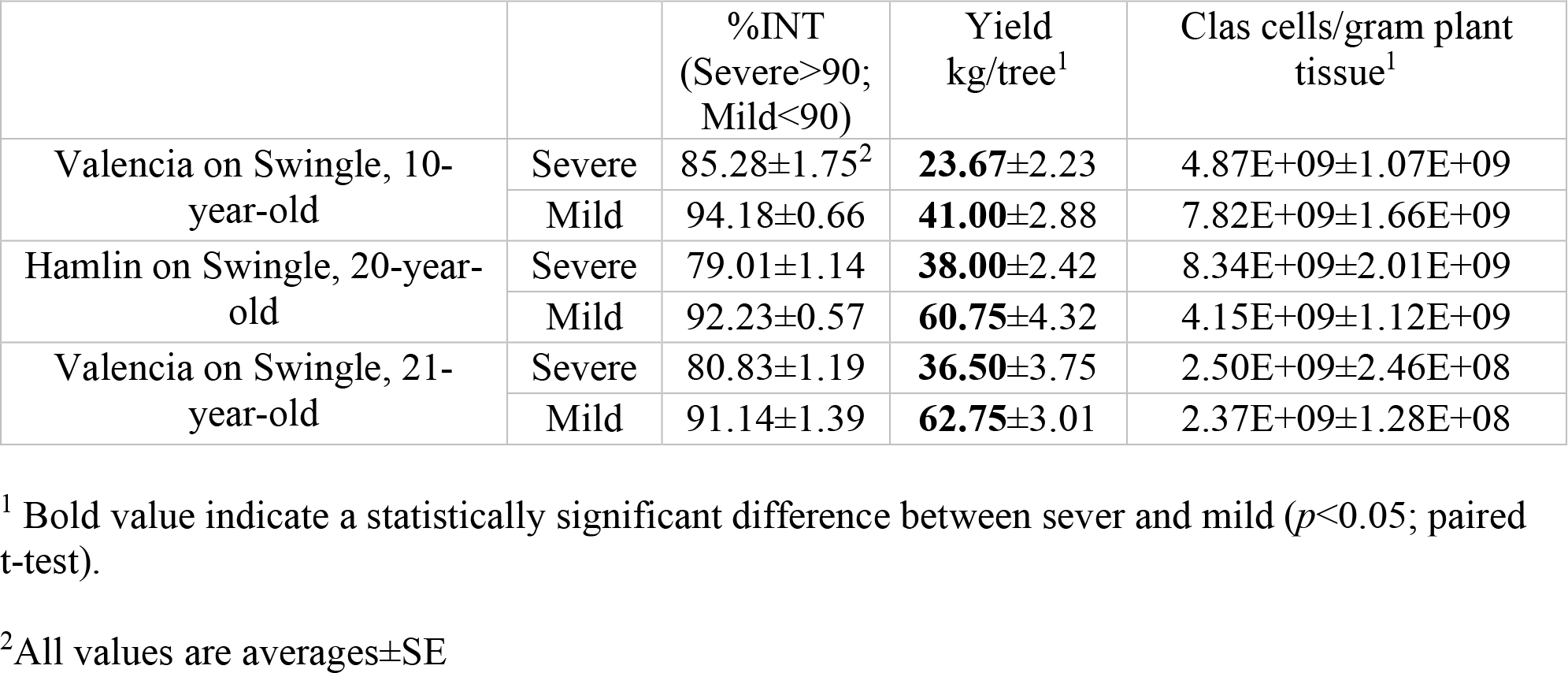
Estimating HLB disease severity according to %INT

We also performed the measurement of macronutrients, which include nitrogen (N), phosphorus (P), potassium (K), calcium (Ca), sulfur (S), magnesium (Mg) and micronutrients, which include boron (B), zinc (Zn), manganese (Mn), iron (Fe), copper (Cu). The results showed there were no significant differences among the trees in the fields (data not shown).

### Effect of photosynthetically active radiation interception in the canopy

We investigated the relationships between yield and %INT from two citrus varieties in three blocks of different fields in Sept. 2019. Total 28 trees were selected for the investigation. A critical examination of data indicated that the yield was positively correlated with %INT. The yield of HLB-affected trees increased as the %INT value increased. Linear regression model was applied to the relationship (Figure 1), and the slope gives a specific value of the increment by %INT.

**Figure 1.**
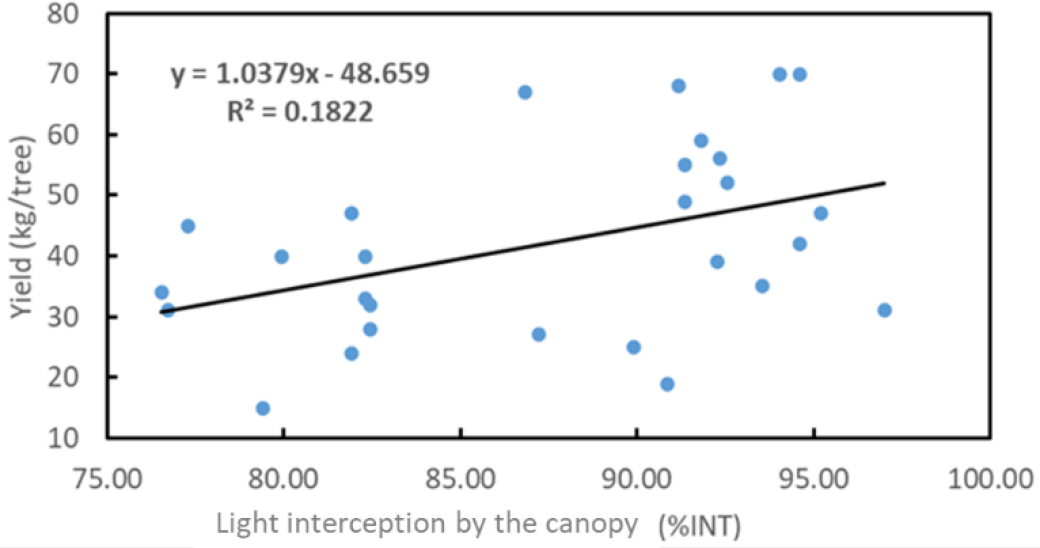
Correlation between yield and photosynthetically active radiation interception percent in the canopy (%INT)

### Effect of FDF and canopy volumes

The perusal of the data (Tables 1) reveals that FDF had significantly impact on the yield (r= 0.468, *p*=0.038). Linear regression model was applied to the relationship between the fruit detachment force and the yield, and the slope gives a specific value of the yield increment by FDF (Figure 2). A similar trend was observed for canopy volume, where canopy volumes were also associated with yields (r=0.542, *p*=0.003) (Table 1), and the linear regression model was applied to the relationship as well (Figure 3).

**Figure 2.**
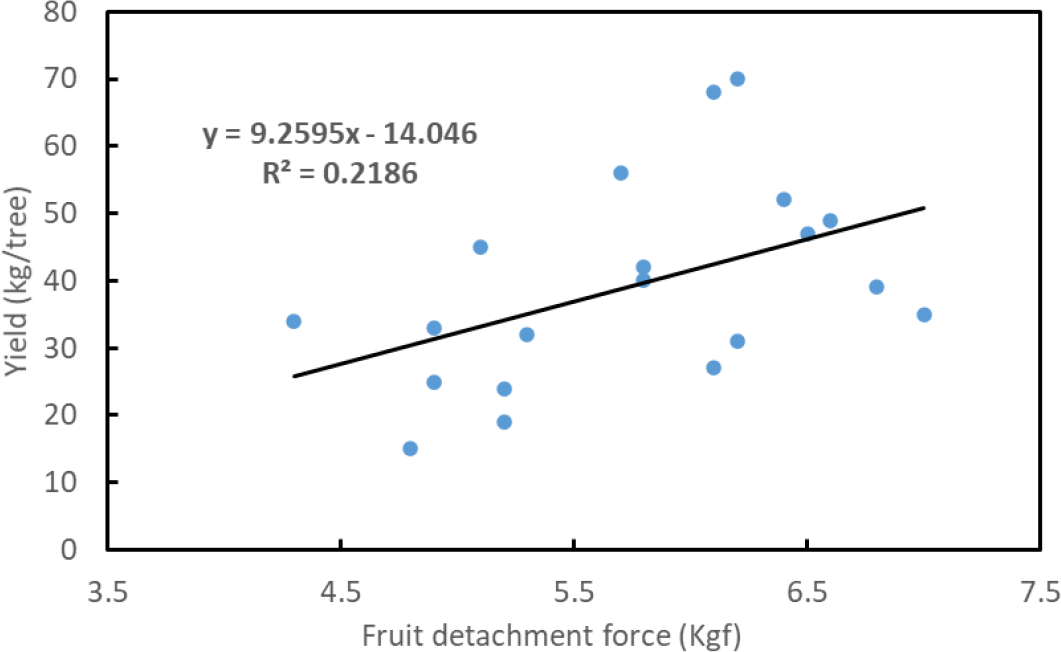
Relationship between the yield and fruit detachment force

**Figure 3.**
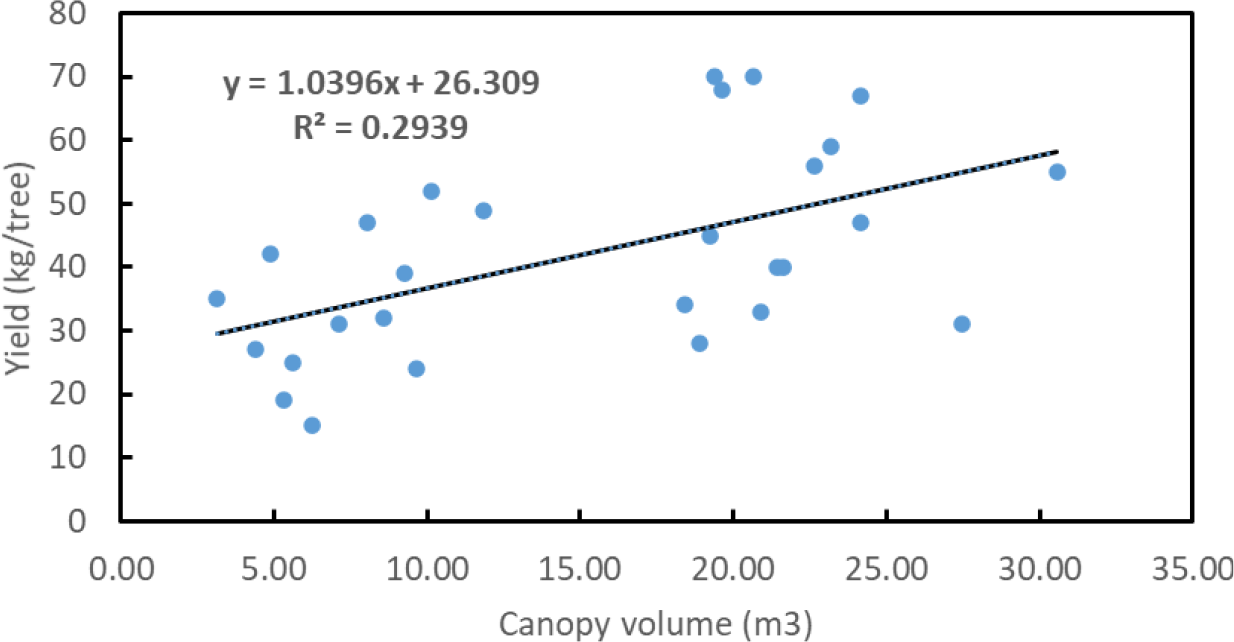
Relationship between the yield and canopy volume

### HLB evaluation in trees according to %INT

Next, we tested if %INT can be used as an accurate measurement to assign disease severity for HLB. Trees were divided into two blocks based on %INT, where trees with an %INT of 90 and over were categorized as mild HLB, and those below 90 were categorized as severe HLB. In all three field trials, trees that were categorized as ‘severe HLB’ according to the %INT had around half of the yield levels compared to those categorized as the ‘mild HLB’ category. Remarkably, there was no difference in amount of CLas cells per gram tissue, as determined by qPCR Ct value. These results indicate that %INT, but not Ct values, can be used accurately to determine HLB disease levels.

### qPCR may not always accurately detect live CLas cells

To test if Ct value is a reliable measure for number of bacteria, we used quantitative PCR and transmission electron microscopy (TEM) in order to visualize CLas in *Citrus macrophilla* (Cmac) and grapefruit plants in greenhouse that were infected with HLB. qPCR and TEM analysis were conducted on the same leaf midribs and results were compared (Table 3). In grapefruit, five months after infection by ACP, in plants that had ‘negative’ Ct values (Ct value above 32) we could not find any CLas cells in our TEM analysis, but we could find few bacterial cells (2 cells in 31 sieve elements examines) in the plant that had a ‘positive’ Ct value (Ct value below 32). In contrast, in Cmac, we detected the highest number of CLas cells (8 CLas cells in 10 sieve elements examined) in the phloem of a plant that was ‘negative’ according to the Ct value. Likewise, in the midrib that had positive Ct value, we could not detect any CLas cells in any of the sieve elements examined (Table 3, Figure 4). These results show that on some occasions qPCR does not reliably detect CLas cells, indicating that the sensitivity of the analysis is not optimal.

**Table 3:** Comparison between *Candidatus* Liberibacter asiaticus quantification by qPCR and TEM

| Plant | Ct value<sup>1</sup> | SE<sup>2</sup> | CLas cells<sup>3</sup> |
| --- | --- | --- | --- |
| **C mac** | 32.09 | 10 | 8 |
| **C mac** | 31.2 | 25 | 0 |
| **C mac** | 32.75 | 14 | 0 |
| **Grapefruit** | 34.9 | 45 | 0 |
| **Grapefruit** | 34.7 | 31 | 0 |
| **Grapefruit** | 28.5 | 31 | 2 |

<sup>1</sup> Ct value <32 is considered positive
<sup>2</sup>Number of sieve elements examined
<sup>3</sup> Number of CLas cells detected

**Figure 4:**
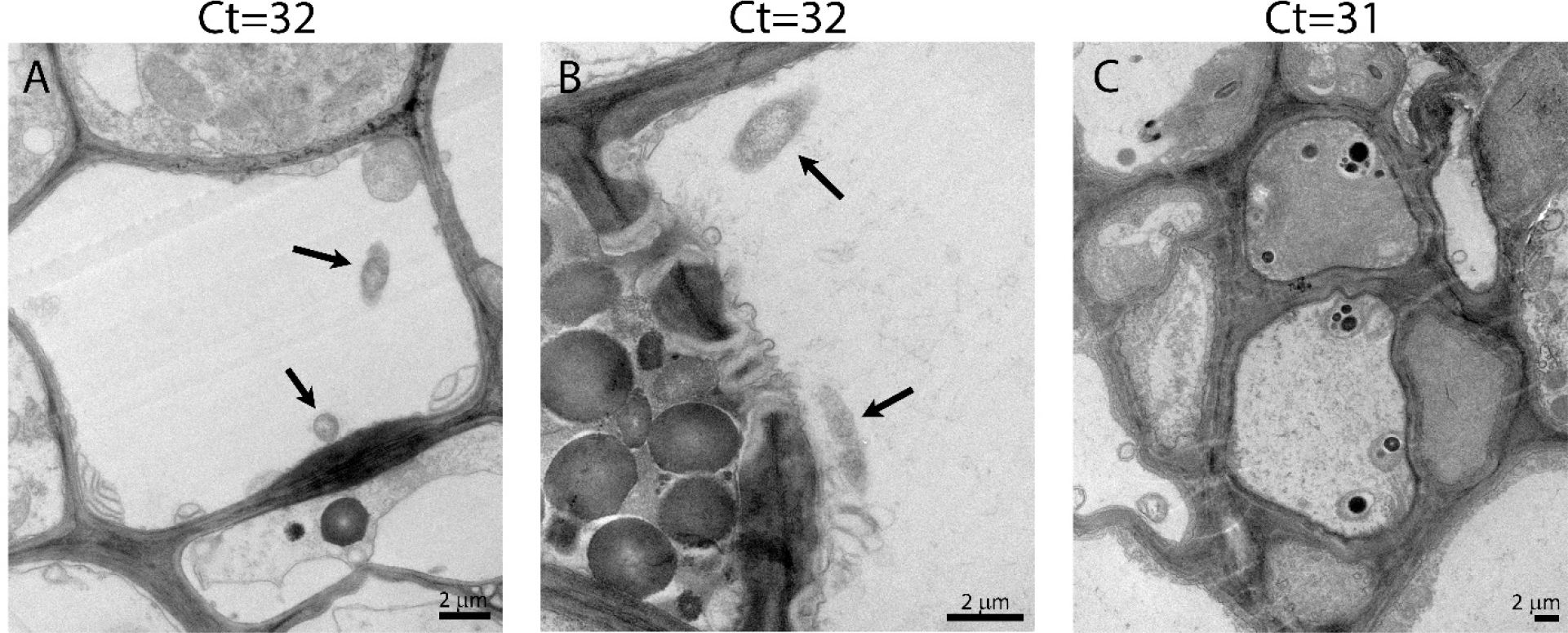
CLas cells in Cmac phloem. CLas bacterial cells were found in midrib sieve elements of a plant with Ct values of 32 (A, B), but no cells were found in midrib of plant with Ct value of 31 (C). CLas cells are marked by an arrow.

## Discussion

Accurate evaluation of the HLB disease status is of major importance to test the efficiency of treatments, especially in field settings. Predicting the number of CLas cells per gram tissue is one of the most common ways of estimating HLB disease status. However, the correlation between CLas levels and disease severity was not proven. Here, the CLas Ct values and numbers of CLas cells were correlated against fruit yield in three different field trials, but no significant correlation was found. These results indicate that Ct value is not a good predictor of HLB disease severity. The lack of correlation may result from a lack of sensitivity of the PCR method, especially as CLas is phloem localized and its levels are low in infected citrus trees [21]. Another possible explanation may be that qPCR is detecting dead, as well as living, bacteria. In fact, Etxeberria et al (2019) [22] measured CLas qPCR Ct values in leaves of ‘Valencia’ orange trees after a heat-treatment that eliminated viable CLas, and found that although titer declined, CLas remained detectable 5 months after treatment, supporting this idea. Here we showed that we could find viable CLas in leaves of Cmac that had a Ct that’s is considered as negative (above 32), suggesting an even more complex situation. Previous work showed there were almost no CLas present in flush tissues of HLB affected plants with Ct values indicative of HLB infection [20]. In that work, it was shown that although no CLas were detected in these leaves, the host plants still responded to the bacterial infection, and the phloem were plugged with callose. Thus, it is also possible that HLB disease is not directly correlated by the number of bacteria in the stem, and the host plant responses may play a significant role in disease development.

We analyzed different possible field measurements for assigning HLB disease severity in field settings and identified three values that were positively correlated with the yield, with very similar correlation coefficient values: the canopy volume, FDF, and %INT. These values are all important parameter in the study of yield estimates in horticulture and are all known to be related to HLB symptomology and are therefore expected to serve as reliable indicators to assess effects therapeutic treatments on diseased plants. For example, it was already shown that HLB affected trees display higher premature fruit drop, and lower FDF [23, 24]. Comparing susceptible HLB varieties to tolerant ones also showed that lower drop rate is associated with HLB tolerance [25]. In HLB tolerant varieties, internal structures were better preserved compared to susceptible ones [26]. It was suggested that CLas effector SDE1 is inducing senescence in citrus, thus providing a possible link between the bacteria and the fruit drop phenomena [27].

Dividing disease severity of trees according to %INT in three independent field trials (representing different varieties and tree ages) into severe or mild, resulted in clear low and high yield categories, respectively. Intercepted PAR (INT) indicates the amount of PAR caught by various canopy layers as the PAR travels through the canopy. % INT was measured above an inside the canopy and represents the amount of light absorbed by the leaves (transmissibility). The %INT value is represented as percentage (of PAR inside the canopy compared to the PAR at the edge), such that higher %INT indicates higher canopy density. Using the %INT has the advantage of being a relative rather than absolute measurement. Therefore, %INT measurement is not dependent on the absolute size of the canopy, or on the specific weather on the day of measurements and could be appropriate for field settings.

Overall, our results show that fruit yield in HLB-affected trees is directly correlated with the the absorption of light by the leaves (higher light absorption results in higher yields), volume of the canopy (bigger canopy gives higher yields), and with the FDF (stronger connection force results in higher yields). Employing these measurements with trees in the field can be used to evaluate HLB severity. We would emphasize that % INT or LAI is a more holistic measurement for HLB-affected trees as it is found to be correlated to fruit size, FDF, and root density. Our study shows that evaluating CLas levels according to Ct values in leaves should not be employed to determine HLB disease levels. It is still not clear if this lack of correlation results from a problem with PCR sensitivity, or uneven distribution of *C*Las in the tree (making sampling difficult), or that the development of HLB does not depend on the level of the bacteria. This lack of connection may indicate that treatments aimed at lowering the bacteria levels in the trees may not have a significant effect on HLB.

## Acknowledgment

The work was funded by the National Institute of Food and Agriculture grant number 2020-70029-33197, and by the Florida State legislative funding for the UF/IFAS Citrus Initiative

